# Deciphering sex-specific genetic architectures using local Bayesian regressions

**DOI:** 10.1101/653386

**Authors:** Scott A Funkhouser, Ana I Vazquez, Juan P Steibel, Catherine W Ernst, Gustavo de los Campos

## Abstract

Many complex human traits exhibit differences between sexes. While numerous factors likely contribute to this phenomenon, growing evidence from genome-wide studies suggest a partial explanation: that males and females from the same population possess differing genetic architectures. Despite this, mapping gene-by-sex (G×S) interactions remains a challenge likely because the magnitude of such an interaction is typically and exceedingly small; traditional genome-wide association techniques may be underpowered to detect such events partly due to the burden of multiple test correction. Here, we developed a local Bayesian regression (LBR) method to estimate sex-specific SNP marker effects after fully accounting for local linkage-disequilibrium (LD) patterns. This enabled us to infer sex-specific effects and G×S interactions either at the single SNP level, or by aggregating the effects of multiple SNPs to make inferences at the level of small LD-based regions. Using simulations in which there was imperfect LD between SNPs and causal variants, we showed that aggregating sex-specific marker effects with LBR provides improved power and resolution to detect G×S interactions over traditional single-SNP-based tests. When using LBR to analyze traits from the UK Biobank, we detected a relatively large G×S interaction impacting bone-mineral density within *ABO* and replicated many previously detected large-magnitude G×S interactions impacting waist-to-hip ratio. We also discovered many new G×S interactions impacting such traits as height and BMI within regions of the genome where both male- and female-specific effects explain a small proportion of phenotypic variance (R^2^ < 1×10^−4^), but are enriched in known expression quantitative trait loci. By combining biobank-level data and techniques to estimate sex-specific SNP effects after accounting for local-LD patterns, we are providing evidence that numerous small-magnitude G×S interactions exist to influence sex differences in a variety of complex traits.

**Author Summary:** Many complex human traits are known to be influenced by an impressive number of causal variants each with very small effects, posing great challenges for genome-wide association studies (GWAS). To add to this challenge, many causal variants may possess context-dependent effects such as effects that are dependent on biological sex. While GWAS are commonly performed using specific methods in which one single nucleotide polymorphism (SNP) at a time is tested for association with a trait, alternatively we utilize methods more commonly observed in the genomic prediction literature. Such methods are advantageous in that they are not burdened by multiple test correction in the same way as traditional GWAS techniques are, and can fully account for linkage-disequilibrium patterns to accurately estimate the true effects of SNP markers. Here we adapt such methods to estimate genetic effects within sexes and provide a powerful means to compare sex-specific genetic effects.

## Introduction

Sex differences are widespread in nature, observed readily among many human traits and diseases. For quantitative traits, sex may affect the distribution of phenotypes at various levels, including mean-differences between genetic males and genetic females (hereafter referred to as males and females, respectively) as well as differences in variance. Sex differences are likely due to a myriad of factors including differential environmental exposures, unequal gene dosages for sex-linked genes as well as sex-heterogeneity in the architecture of genetic effects at one or more autosomal loci (i.e. gene-by-sex (G×S) interactions). In this way, sex is considered an environmental variable, providing two well-defined conditions in which allele frequencies and linkage disequilibrium (LD) patterns are equivalent but nevertheless genetic effects of one or many autosomal loci may differ.

Evidence for different genetic architectures between sexes among human populations is largely supported by genome-wide parameters [1–4] including unequal within-sex heritabilities (*h*^2^_male_ ≠ *h*^2^_female_) and between-sex genetic correlations less than one (*r*_g_ < 1); the former suggests that the proportion of phenotypic variance explained by genetic factors varies between sexes, while the latter suggests genetic effects are disproportional between sexes [5]. Although many traits seem to have between-sex genetic correlation that is evidentially less than one, genome-wide association (GWA) studies intended to map G×S interactions have struggled to pinpoint such loci [6,7]. Based on this dichotomy, G×S interactions presumably exist for many traits, but the magnitude of a typical G×S interaction is suspected to be exceedingly small, explaining why such events commonly elude detection, particularly after multiple test correction. However, just as numerous small effect causal loci accumulate to affect phenotypic variance, small G×S interactions may accumulate to influence both sex differences and phenotypic variance.

Most GWA studies utilize single-marker regression (SMR), in which the phenotype is regressed upon allele content one SNP at a time, thereby obtaining marginal SNP effect size estimates that do not fully account for LD patterns. In contrast, whole-genome regression methods, in which the phenotype is regressed upon all SNPs across the genome concurrently, fully account for multi-locus LD. These methods are increasingly being used as a one-stop solution to estimate true (conditional) effect sizes of SNP markers and to provide genome-wide estimates including genomic heritability [8–10] and between-sex genetic correlations [2–4]. By estimating true SNP effect sizes, the goal across many studies is to select SNPs with non-zero effects and to build a model for predicting polygenic scores [11–13]. Other works have directly illustrated the use of whole-genome regression methods for GWAS [14–17]. Whole-genome regressions are computationally challenging to use with biobank-level data; however, recent work suggests relatively accurate genomic prediction and SNP effect estimation can be achieved by simply accounting for local LD patterns (as opposed to global LD patterns) [18].

Building on the idea of utilizing true SNP marker effects, here we developed local Bayesian regressions (LBR) in which the phenotype is regressed upon multiple SNPs spanning multiple LD blocks (thereby accounting for local LD patterns) to study sex differences in complex traits from the UK Biobank. The LBR model uses random-effect SNP-by-sex interactions [19,20] that decompose conditional SNP effects into three components: i) one shared across sexes, ii) a male-specific deviation from the shared component, and iii) a female-specific deviation from the shared component. Using samples from the posterior distribution of conditional SNP effects, we developed methods to infer sex-specific effects and G× S interactions at the single SNP level and by aggregating SNP effects within small LD-based regions, offering multiple perspectives to study sex-specific genetic architectures.

In this study, we have utilized genotypes for 607,497 autosomal SNPs from ~259,000 distantly related Caucasians from the UK Biobank for assessing LBR’s performance in analyzing simulated and real complex traits including height, BMI, waist-to-hip ratio (WHR), and heel bone-mineral density (BMD). Simulations showed that (i) for inferences of G×S interactions, LBR offers higher power with lower FDR than methods based on marginal effects (aka single-marker regression) and (ii) we show that under imperfect LD between SNPs and causal variants (i.e., when causal variants are not genotyped), aggregating SNP effects within small LD-based regions offers higher power than methods based on testing individual SNPs.

The traits analyzed in this study span a range of genome-wide metrics and G×S suggestibility; from height and BMI for which previous studies indicate males and females possess very similar genetic architectures [3], to WHR, a trait with well-documented G×S interactions [21–24], and BMD, for which G×S interactions are thought to exist but have eluded detection [25]. LBR provided evidence of G×S interactions impacting height, BMI, and BMD at regions of the genome where sex-specific genetic effects are relatively small, however such regions are enriched in known eQTL. For WHR, LBR replicated many large-magnitude G×S interactions previously discovered using single-marker regression, but also located novel G×S interactions near such genes as the estrogen receptor *ESR1*.

## Results

### Overview of the LBR model, inference methods, and implementation

To study sex differences we regressed male and female phenotypes (***y***_m_ and ***y***_m_) on male and female genotypes (***X***_m_ and ***X***_f_) using a SNP-by-sex interaction model of the form

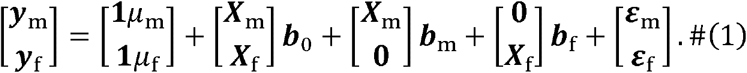

Above, *μ*_m_ and *μ*_f_ are male and female intercepts, ***b***_0_ = {*b*_0_*j*__} (*j* = 1,…, *p*) is a vector of main effects, ***b***_m_ = {*b*_m_*j*__} and ***b*_f_** = {*b*_f_*j*__} are male and female interactions, respectively and ***ε***_m_ = {*ε*_m_*i*__} and ***ε*_f_** = {*ε*_f_*i*__} are male and female errors which were assumed to follow normal distributions with zero mean and sex-specific variances. Female-specific and male-specific SNP effects are defined as *β*_f_*j*__ = *b*_0_*j*__ + *b*_f_*j*__ and *β*_m_*j*__ = *b*_0_*j*__ + *b*_m_*j*__, respectively.

#### Prior assumptions

For SNP effects we adopted priors from the spike-slab family with a point of mass at zero and a Gaussian slab [26] specifically, 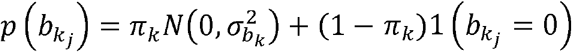 (where *k* = 0, f or m). Here, *π_k_* and 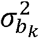 are hyper-parameters representing the proportion of nonzero effects and the variance of the slab; these hyper-parameters were treated as unknown and given their own hyperpriors (see Methods).

#### Local-regression

Implementing the above model with whole-genome SNPs (*p* ~ 600K and very large sample size (*n* ~ 300K) is computationally extremely challenging. However, LD in homogeneous un-structured human populations spans over relatively short regions (R^2^ between allele dosages typically vanishes within 1-2 Mb; S1 Fig). Therefore, we applied LBR to long, overlapping chromosome segments (Fig. 1). Specifically, we divided the genome into “core” segments containing 1,500 contiguous SNPs (roughly 8Mb, on average), then applied the regression in equation (1) to SNPs in the core segment plus 250 SNPs (i.e., roughly 1Mb) in each flanking region, which were added to account for LD between SNPs at the edge of each core segment with SNPs in neighboring segments.

**Fig 1.**
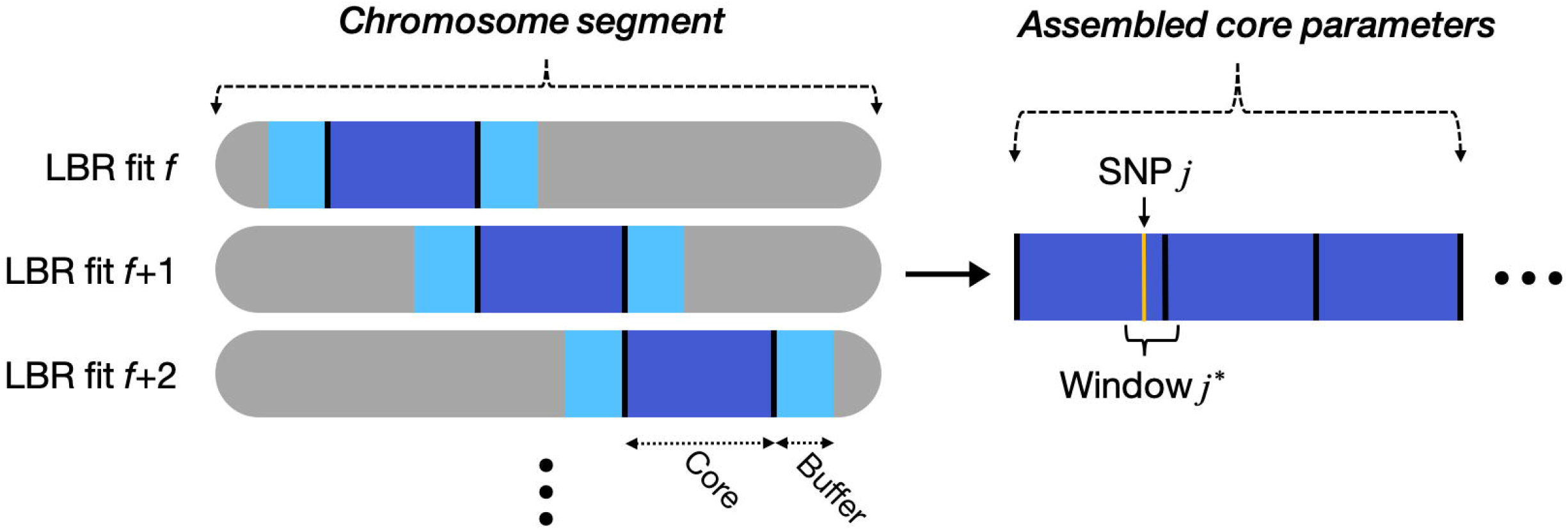
Strategy for implementing local Bayesian regressions genome-wide. The phenotype is regressed upon multiple sequential SNPs using a sliding window approach. The core region contained 1500 SNPs (roughly 8Mb, on average) and each buffer region contained 250 SNPs (roughly 1Mb, on average). Core parameters (posterior samples) are stitched together, then sex-specific effects and G×S interactions are inferred at the level of SNP *j* and window *j*\*.

#### Inferences

We used the BGLR [27] software to draw samples from the posterior distribution of the model parameters and used these samples to make inference about individual SNP effects including: (i) the posterior probability that the *j*^th^ SNP has a nonzero effect in males (PPM_SNP_*j*__) and females (PPF_SNP_*j*__) and (ii) the posterior probability that the female and male effects are different(PPDiff_SNP_*j*__).

In regions involving multiple SNPs in strong LD, inferences at the individual-SNP level may be questionable. Therefore we borrowed upon previous work by Fernando et al. [14], enabling us to aggregate multiple sex-specific SNP effects within relatively small regions using “window variances”. For each SNP *j* we defined a window *j*\* around the SNP based on local LD patterns (see Methods). We then defined the male-specific and female-specific window variances as 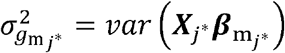 and 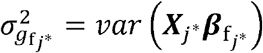, respectively. Here, ***X***_*j*\*_ represent genotypes at SNPs within the *j*\* window and *var*() is the sample variance operator. Prior to model fitting, the phenotype is scaled across sexes; thus, sex-specific window variances may be interpreted as the proportion of total phenotypic variance explained by sex-specific SNP effects. From samples of sex-specific window variances, we computed the posterior probability of (i) nonzero male-specific window variance 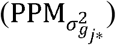, (ii) nonzero female-specific window variance 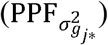, and (iii) sex difference in window variances (denoted as 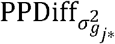).

### Local Bayesian regressions offer improved power with lower false-discovery rates

We used simulations to assess the power and false discovery rate (FDR) of LBR and to compare it with that of standard single-marker-regression (SMR). Traits were simulated using SNP genotypes from the Axiom UK-Biobank (119,190 males and 139,738 females, all distantly related Caucasians). We simulated a highly complex trait with one causal variant (CV) per ~2Mb which on average explained a proportion of the phenotypic variance equal to 3.3×10^−4^. Our simulation used a total of 60,000 SNPs (consisting of 6,000 consecutive SNPs taken from 10 different chromosomes) and 150 CVs; on the complete human genome “scale” this corresponds to a trait with 1,500 CVs and a heritability of 0.5 (see Methods for further details). 40% of the CVs (a total of 60 SNPs in our simulation) had differing sex-specific effects and the remaining 60% (90 SNPs) had effects that were the same in males and females.

#### Power and FDR when causal variants are genotyped

First, we analyzed the simulated phenotypes using all SNPs (including the 150 causal ones). Initially interested in inferring G×S interactions, we ranked SNPs based on LBR’s PPDiff_SNP_*j*__. metric and based on SMR’s *p*-value for sex difference (*pvalue*-diff, see Methods) and used the two ranks to estimate power and FDR as a function of the number of SNPs selected (Fig 2). LBR showed consistently higher power (achieving a power of ~80% when selecting the top 50 SNPs with highest PPDiff_SNP_*j*__) and lower FDR than SMR. The false discovery rate of LBR was very low when selecting the top-50 SNPs with highest PPDiff_SNP_*j*__ and exhibited a very sharp phase-transition with fast increase in FDR thereafter.

**Fig 2.**
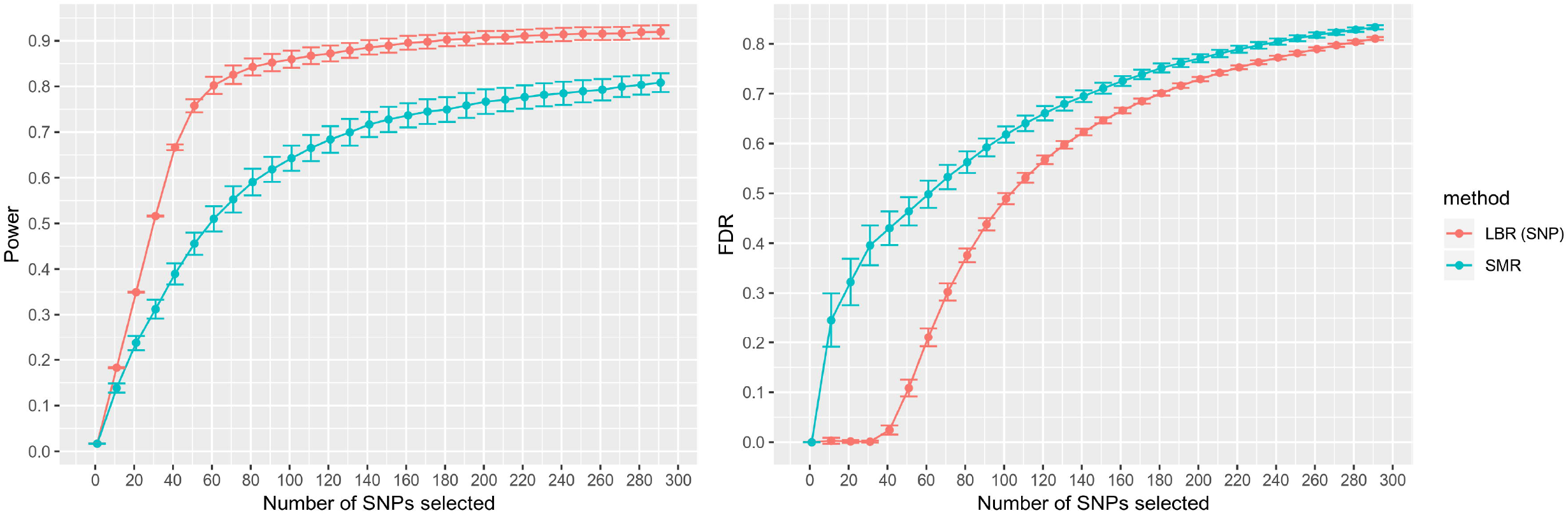
Estimated power and false-discovery rate for discovering observed SNPs with G×S interactions. Shown as a function of the number of SNPs selected. Each point represents a sample average and error bars represent 95% confidence intervals, each derived using 30 Monte Carlo replicates. LBR (SNP): local Bayesian regression, utilizing PPDiff_SNP_*j*__ SMR: single-marker regression, utilizing *pvalue*-diff.

We also compared the two methods based on arbitrary, albeit commonly used, mapping thresholds for SMR and LBR. At PPDiff_SNP_*j*__ ≥ 0.95, LBR selected an average (across simulation replicates) of 38.33 SNPs with an estimated power of 0.634 and estimated FDR of 0.007. Conversely, at *pvalue*-diff ≤ 5×10^−8^, SMR selected an average of 50.7 SNPs with an estimated power of 0.436 and estimated FDR of 0.451. Altogether, these results suggest that for G× S discovery, LBR offers higher power and lower FDR than SMR—the method most widely used in GWA studies—at least when G×S interactions are observed.

When trying to map SNPs that had effect in at least one sex, we used PP_SNP_*j*__ = max[PPM_SNP_*j*__, PPF_SNP_*j*__] and *p*-values from an F-test (see Methods) as metrics for LBR and SMR methods, respectively. Again, LBR showed higher power with lower FDR than a standard SMR *p*-value (S1 Fig). At traditional mapping thresholds, LBR and SMR had similar power but LBR achieved that power with much lower FDR; at PP_SNP_*j*__ ≥ 0.95, the average number of SNPs selected was 120.83 with an estimated power of 0.799 and estimated FDR of 0.009 while at *p*-value ≤ 5×10^−8^, the number of SNPs selected was 374.56 with an estimated power of 0.794 and FDR of 0.66.

#### Power and FDR under imperfect LD

In a second round of analyses, we removed all CVs from the panel of SNPs used in the analysis to represent a situation where CVs are not observed, and genotyped SNPs are tagging CVs at varying degrees. As before, we initially assessed the relative performance of LBR to infer segments harboring G×S interactions. Power and FDR were assessed at several resolutions: 1Mb, 500Kb and 250Kb regions around each CV. At each resolution, a discovery was considered true if the finding laid within a segment harboring a G×S CV. Power and FDR were computed at different thresholds (PPDiff_SNP_*j*__ and 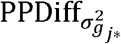 for LBR and *pvalue*-diff for SMR; Fig 3). When using a 1Mb target area—such that correct G×S discoveries must be within 500Kb on either side of a true G×S event—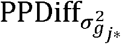 thresholds (LBR’s window-based metrics) provided highest power within an FDR range of 0-0.3, thereafter SMR provided slightly higher power. As expected, when removing CVs, power was estimated to be much lower than when CVs were observed; at 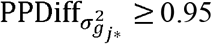, the estimated power and FDR were 0.454 and 0.004, respectively, while at *pvalue*-diff ≤ 5×10^−8^, estimated power and FDR were 0.22 and 0.006. As seen in Fig 3, when considering a finer resolution (500Kb and 250Kb) the performance of both LBR-based approaches was more robust than SMR. Altogether this indicates that for the discovery and mapping of unobserved G×S interactions, LBR’s window-based metric provides higher power with equivalent FDR and finer resolution than single-marker regression methods.

**Fig 3.**
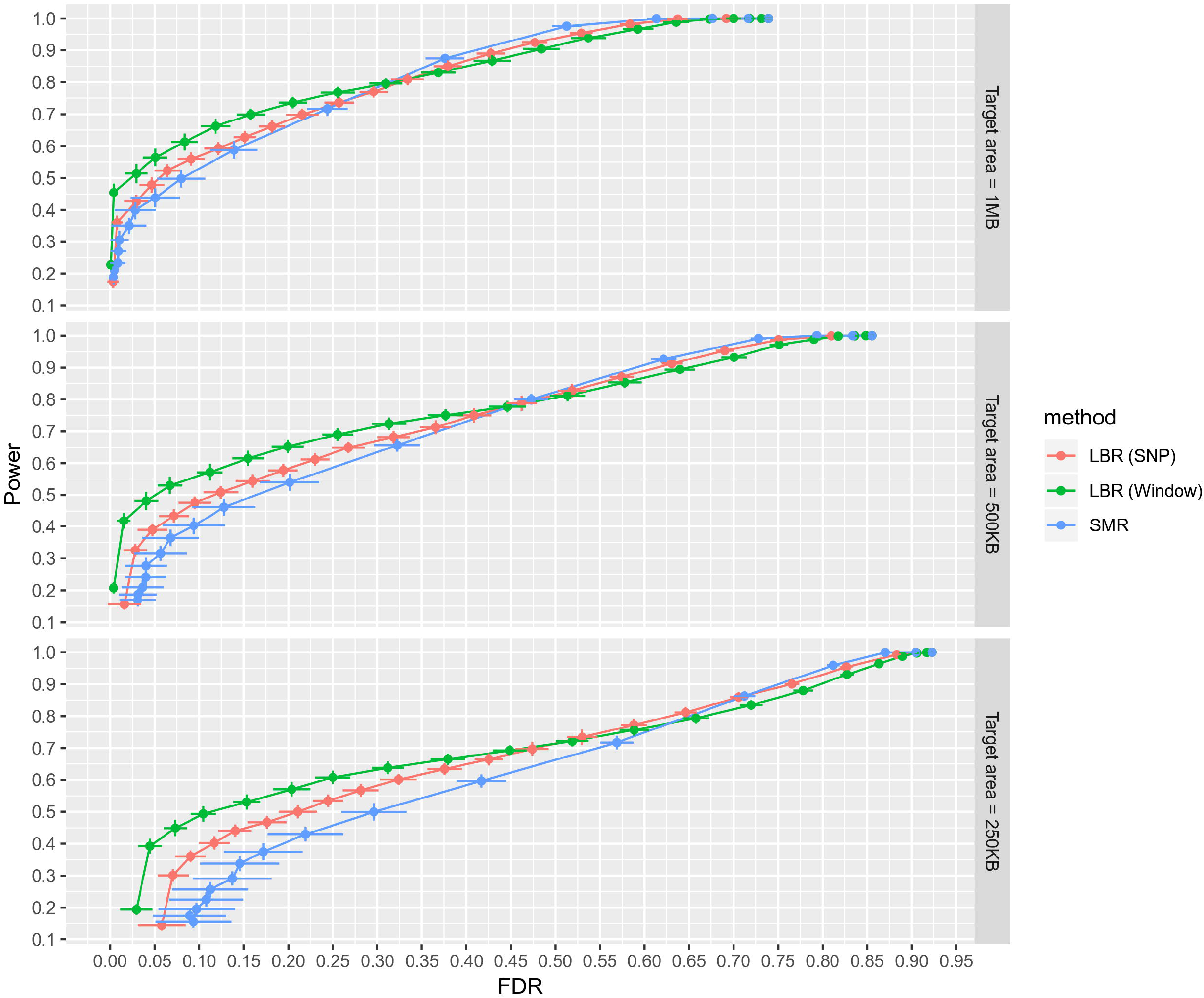
Power vs false-discovery rate for discovering genomic regions containing masked G×S interactions. Here power is defined as the expected proportion of G×S interactions that are being tagged by at least one selected SNP *j* or window *j*\*. False discovery rate is defined as the expected proportion of selected SNPs or windows that are not tagging any G×S interactions. Each point is an estimate and error bars for both axes represent 95% confidence intervals. Point estimates and intervals were derived using 30 Monte Carlo replicates. Each facet corresponds to a different “target area”, a fixed width around each G×S interaction that defines the set of SNPs effectively tagging it. LBR (SNP): uses the PPDiff_SNP_*j*__ metric spanning 1-0. LBR (Window): uses the 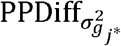 metric spanning 1-0. SMR: uses the *pvalue*-diff metric spanning (on the −log_10_ scale) 8-0.

To infer segments containing CVs that affect at least one sex, we again used LBR to decide whether either sex-specific effect was nonzero at the level of individual SNPs or windows. Using a 1MB target area, LBR’s window-based metrics provided the highest power within an FDR range of 0-0.025. When decreasing the target area, LBR provided the highest power over larger FDR ranges (S2 Fig).

### For real human traits, many newly discovered G×S interactions show relatively small sex-specific effects

We analyzed four complex human traits (height, BMI, BMD, and WHR) measured among ~259,000 distantly related Caucasians from the UK Biobank (~119,000 males and ~140,000 females). For each trait, we fit the LBR model (eq. 1) across the entire autosome consisting of 607,497 genotyped SNPs using 417 overlapping segments (Fig. 1) and obtained evidence of G×S interactions at the level of SNP *j* and window *j*\*.

To compare both the magnitude and sign of sex-specific SNP effects, we plotted each 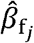 against 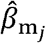 (Fig 4a). The trait was scaled across sexes prior to model fitting; thus, male- and female-specific effects were not constrained to the same scale. In this way, one might expect male-specific SNP effects to uniformly differ from female-specific SNP effects by a multiplicative factor if the variance of the phenotype is different between sexes (sample statistics within each sex are provided within S1 Table). Surprisingly, we did not observe evidence of sex-specific SNP effects uniformly differing due to differences in phenotypic scale; for height, BMD, and BMI, as seen in Fig. 4a, most large effect SNPs lie near the blue diagonal line. For WHR, we observed largely consistent results from prior studies [21–23]: namely the prevalence of numerous SNPs with relatively large effects in females but little to no effect in males. No traits exhibited evidence of any SNPs with (i) high confidence male- and female-specific effects (no SNPs with PPM_SNP_*j*__ ≥ 0.9 and PPM_SNP_*j*__ ≥ 0.9) and (ii) differing signs between sexes.

**Fig 4.**
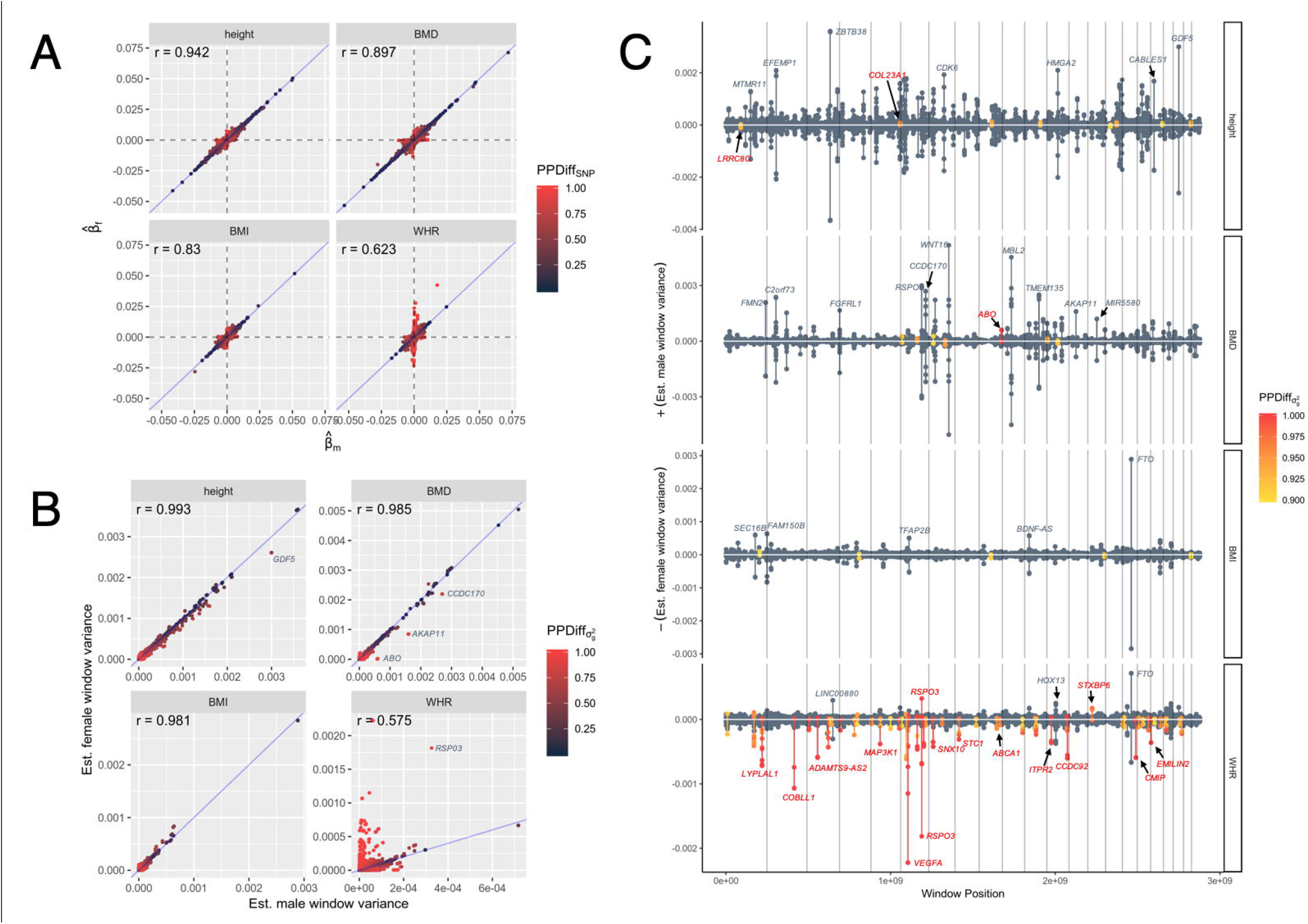
Comparing sex-specific genetic effects. **(A)** Plot of estimated female SNP effects against estimated male SNP effects for all 607,497 genotyped autosomal SNPs. Points are colored by their posterior probability of sex difference at the level of individual SNPs. **(B)** Plot of estimated female window variances against estimated male window variances for all 607,497 LD-based windows, with each window *j*\* centered on a different focal SNP *j*. Points are colored by their posterior probability of sex difference at the level of window variances. **(C)** Miami-like plot depicting location and magnitude of G×S interactions identified through sex-specific window variances. For each trait, showing estimated male window variance above the x-axis and estimated female window variance below the x-axis. Vertical lines denote changing chromosomes. A sample of windows is labeled with nearest gene annotation, obtained from Axiom UKB WCSG annotations, release 34. Gray labels indicate nearest genes with relatively large window variances evidently shared across sexes, while red labels indicate nearest genes with detected G×S interactions.

We then aggregated sex-specific SNP effects within small LD-based regions to estimate sex-specific window variances 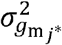 and 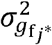 and compared the magnitude of each (Fig 4b). Interestingly for traits such as height, many large effect regions bear slightly larger window variances for males than for females. This was not observed at the single SNP level, suggesting that many regions bearing numerous small effect SNPs produce aggregate effects that are potentially larger (although not reaching a 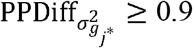 threshold) in males than in females.

One example is the GDF5 locus, previously known to strongly associate with adult height [28], where a peak 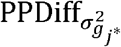 signal centered on rs143384 had slightly different estimated sex-specific window variances (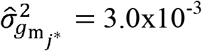 and 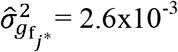) but weak evidence of a G×S interaction 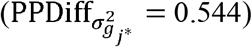. For BMD, several large effect regions show suggestive evidence of G×S interactions including the *AKAP11* locus and the *CCDC170* locus (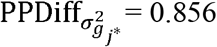 and 0.745, respectively), both previously associated with bone mineral density [29–32].

To make G×S inferences at the level of window variances irrespective of the magnitude of sex-specific effects, we adopted a 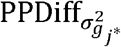 threshold of 0.9, which in simulations (Fig 3) provided optimal power at an estimated FDR of 0.029 when using a 1MB target area. For height, a total of eight distinct regions possessed a 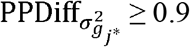, two of which possessed a 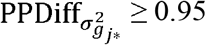. For BMI, 5 distinct regions possessed a 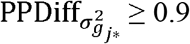 with none reaching a more stringent 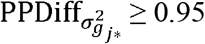 threshold, and none overlapping with two previously suggested BMI G×S SNPs [33]. As seen in Fig 4C, inferred G×S interactions for height and BMI possess relatively small sex-specific window variances; as an example, for height, the window centered on SNP rs1535515 (near *LRRC8C*) had a 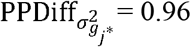, while 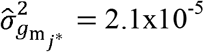 and 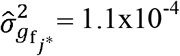. For BMD, seven regions reached a 0.9 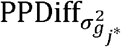 threshold while one higher-confidence G×S interaction 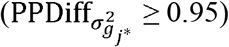 was detected within *ABO*, the gene controlling blood type.

For WHR, roughly 45 distinct genomic regions possessed a 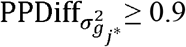, while 34 of these possessed a 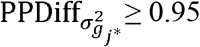. We found many previously detected G×S interactions known to associate with WHR or a related trait, WHR adjusted for BMI (WHRadjBMI) [21–24]. These included interactions at *LYPLAL1, MAP3K1, COBLL1, RSPO3*, and *VEGFA* among others. We also detected numerous novel G×S interactions (Table 1) near physiologically intriguing genes such as the estrogen receptor gene *ESR1* and the ATP binding cassette transporter A1 gene *ABCA1* known to play a role in HDL metabolism 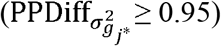. As seen in Table 1, both novel signals possessed a high-confidence female-specific effect with weak evidence for a male-specific effect 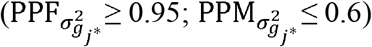, however the magnitude of the female-specific effect was relatively small 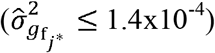. As evident from Table 1, most novel WHR G×S interactions detectable with LBR are those with relatively small sex-specific effects.

**Table 1.**
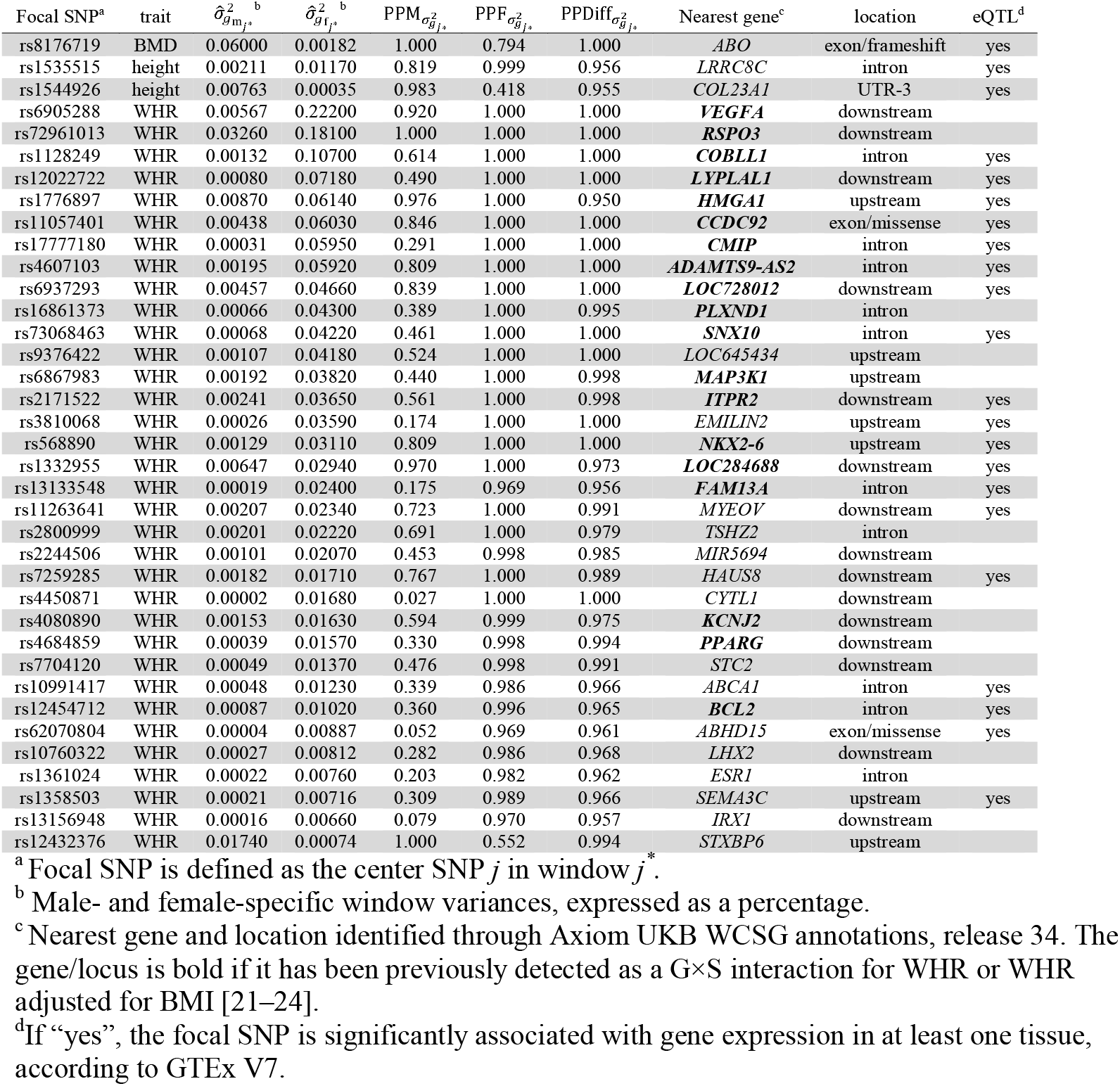
G×S interactions inferred through sex-specific window variances. Listed are loci with at least 0.95 posterior probability that sex-specific window variances differ. The table is sorted first by trait, then by magnitude of the female-specific window variance. Results are filtered such that each window listed consisted of a distinct set of SNPs. A full list of all G×S signals at a 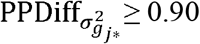 threshold is provided in S2 Table.

| Focal SNP<sup>a</sup> | trait | $\hat{\sigma}_{g_m,j}^2$<sup>b</sup> | $\hat{\sigma}_{g_f,j}^2$<sup>b</sup> | PPM$\sigma_{g,j}^2$ | PPF$\sigma_{g,j}^2$ | PPDiff$\sigma_{g,j}^2$ | Nearest gene<sup>c</sup> | location | eQTL<sup>d</sup> |
| --- | --- | --- | --- | --- | --- | --- | --- | --- | --- |
| rs8176719 | BMD | 0.06000 | 0.00182 | 1.000 | 0.794 | 1.000 | *ABO* | exon/frameshift | yes |
| rs1535515 | height | 0.00211 | 0.01170 | 0.819 | 0.999 | 0.956 | *LRRC8C* | intron | yes |
| rs1544926 | height | 0.00763 | 0.00035 | 0.983 | 0.418 | 0.955 | *COL23A1* | UTR-3 | yes |
| rs6905288 | WHR | 0.00567 | 0.22200 | 0.920 | 1.000 | 1.000 | *VEGFA* | downstream |  |
| rs72961013 | WHR | 0.03260 | 0.18100 | 1.000 | 1.000 | 1.000 | *RSPO3* | downstream |  |
| rs1128249 | WHR | 0.00132 | 0.10700 | 0.614 | 1.000 | 1.000 | *COBLL1* | intron | yes |
| rs12022722 | WHR | 0.00080 | 0.07180 | 0.490 | 1.000 | 1.000 | *LYPLAL1* | downstream | yes |
| rs1776897 | WHR | 0.00870 | 0.06140 | 0.976 | 1.000 | 0.950 | *HMGA1* | upstream | yes |
| rs11057401 | WHR | 0.00438 | 0.06030 | 0.846 | 1.000 | 1.000 | *CCDC92* | exon/missense | yes |
| rs17777180 | WHR | 0.00031 | 0.05950 | 0.291 | 1.000 | 1.000 | *CMIP* | intron | yes |
| rs4607103 | WHR | 0.00195 | 0.05920 | 0.809 | 1.000 | 1.000 | *ADAMTS9-AS2* | intron | yes |
| rs6937293 | WHR | 0.00457 | 0.04660 | 0.839 | 1.000 | 1.000 | *LOC728012* | downstream | yes |
| rs16861373 | WHR | 0.00066 | 0.04300 | 0.389 | 1.000 | 0.995 | *PLXND1* | intron |  |
| rs73068463 | WHR | 0.00068 | 0.04220 | 0.461 | 1.000 | 1.000 | *SNX10* | intron | yes |
| rs9376422 | WHR | 0.00107 | 0.04180 | 0.524 | 1.000 | 1.000 | *LOC645434* | upstream |  |
| rs6867983 | WHR | 0.00192 | 0.03820 | 0.440 | 1.000 | 0.998 | *MAP3K1* | upstream |  |
| rs2171522 | WHR | 0.00241 | 0.03650 | 0.561 | 1.000 | 0.998 | *ITPR2* | downstream | yes |
| rs3810068 | WHR | 0.00026 | 0.03590 | 0.174 | 1.000 | 1.000 | *EMILIN2* | upstream | yes |
| rs568890 | WHR | 0.00129 | 0.03110 | 0.809 | 1.000 | 1.000 | *NKX2-6* | upstream | yes |
| rs1332955 | WHR | 0.00647 | 0.02940 | 0.970 | 1.000 | 0.973 | *LOC284688* | downstream | yes |
| rs13133548 | WHR | 0.00019 | 0.02400 | 0.175 | 0.969 | 0.956 | *FAM13A* | intron | yes |
| rs11263641 | WHR | 0.00207 | 0.02340 | 0.723 | 1.000 | 0.991 | *MYEOV* | downstream | yes |
| rs2800999 | WHR | 0.00201 | 0.02220 | 0.691 | 1.000 | 0.979 | *TSHZ2* | intron |  |
| rs2244506 | WHR | 0.00101 | 0.02070 | 0.453 | 0.998 | 0.985 | *MIR5694* | downstream |  |
| rs7259285 | WHR | 0.00182 | 0.01710 | 0.767 | 1.000 | 0.989 | *HAUS8* | downstream | yes |
| rs4450871 | WHR | 0.00002 | 0.01680 | 0.027 | 1.000 | 1.000 | *CYTL1* | downstream |  |
| rs4080890 | WHR | 0.00153 | 0.01630 | 0.594 | 0.999 | 0.975 | *KCNJ2* | downstream |  |
| rs4684859 | WHR | 0.00039 | 0.01570 | 0.330 | 0.998 | 0.994 | *PPARG* | downstream |  |
| rs7704120 | WHR | 0.00049 | 0.01370 | 0.476 | 0.998 | 0.991 | *STC2* | downstream |  |
| rs10991417 | WHR | 0.00048 | 0.01230 | 0.339 | 0.986 | 0.966 | *ABCA1* | intron | yes |
| rs12454712 | WHR | 0.00087 | 0.01020 | 0.360 | 0.996 | 0.965 | *BCL2* | intron | yes |
| rs62070804 | WHR | 0.00004 | 0.00887 | 0.052 | 0.969 | 0.961 | *ABHD15* | exon/missense | yes |
| rs10760322 | WHR | 0.00027 | 0.00812 | 0.282 | 0.986 | 0.968 | *LHX2* | downstream |  |
| rs1361024 | WHR | 0.00022 | 0.00760 | 0.203 | 0.982 | 0.962 | *ESR1* | intron |  |
| rs1358503 | WHR | 0.00021 | 0.00716 | 0.309 | 0.989 | 0.966 | *SEMA3C* | upstream | yes |
| rs13156948 | WHR | 0.00016 | 0.00660 | 0.079 | 0.970 | 0.957 | *IRX1* | downstream |  |
| rs12432376 | WHR | 0.01740 | 0.00074 | 1.000 | 0.552 | 0.994 | *STXBP6* | upstream |  |

<sup>a</sup> Focal SNP is defined as the center SNP $j$ in window $j^*$ .
<sup>b</sup> Male- and female-specific window variances, expressed as a percentage.
<sup>c</sup> Nearest gene and location identified through Axiom UKB WCSG annotations, release 34. The gene/locus is bold if it has been previously detected as a G×S interaction for WHR or WHR adjusted for BMI [21–24].
<sup>d</sup> If “yes”, the focal SNP is significantly associated with gene expression in at least one tissue, according to GTEx V7.

Additionally, we utilized a traditional SMR approach (see Methods) for the discovery of G×S interactions among traits to compare *pvalue*-diff signals to 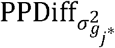 signals (S3 Fig). At *pvalue*-diff ≤ 5×10^−8^, there were no genome-wide significant G×S-interacting SNPs for height, one significant SNP for BMI nearby a window with 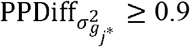, and one significant peak within *ABO* for BMD (the same signal detected using 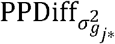). Regions with a 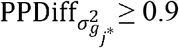 generally coincided with at least nominally-significant *pvalue*-diff signals; for height and BMD, regions with 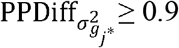 also possessed a peak SNP with *pvalue*-diff ≤ 0.01. For BMI, 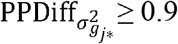 signals possessed a peak SNP of *pvalue*-diff ≤ 0.1. This, together with the fact that novel G×S interactions found using LBR possess relatively small sex-specific effects, suggests that LBR may be detecting G×S interactions that are otherwise missed due to low power. Lastly for WHR, most of the high-confidence 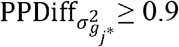 signals coincided with clear and obvious *pvalue*-diff peaks.

### Inferred G×S interactions are enriched in tissue-specific eQTL

As seen previously, many G×S interactions inferred using LBR have exceedingly small sex-specific effects. To further investigate whether G×S detections using the 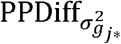 metric may be functionally relevant, we inferred whether such signals are enriched in eQTL identified from GTEx. Specifically, using a hypergeometric test we asked whether 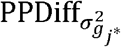-selected focal SNPs (SNP *j* within window *j*\*) were enriched in eQTL, then compared to eQTL enrichment from *pvalue*-diff-selected SNPs as a function of the number of SNPs selected (S4 Fig). 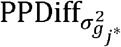-selected focal SNPs showed consistently higher eQTL enrichment than *pvalue*-diff-selected SNPs for all traits except WHR. For instance, at 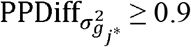, the total number of windows (focal SNPs) selected was 36, 264, 34, and 13, for height, WHR, BMD, and BMI, respectively. With these selections, eQTL enrichment *p*-values were 2.39×10^−4^, 1.52×10^−12^, 2.01×10^−12^, and 8.33×10^−4^, for height, WHR, BMD, and BMI, respectively. When selecting the same number of SNPs using *pvalue*-diff, enrichment *p*-values were 2.25×10^−2^, 1.56×10^−28^, 5.54×10^−8^, 1.93^−1^, for height, WHR, BMD, and BMI, respectively.

To provide more information about how genetic regions bearing G×S interactions may impact gene expression in specific tissues, we determined whether focal SNPs at 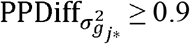 are enriched in tissue-specific eQTL (Fig. 5). For height, BMD, and WHR, such SNPs showed significant eQTL enrichment in at least one tissue, using a conservative bonferroni corrected enrichment *p*-value of 2.6×10^−4^ (correcting for 192 tests in total; 48 tissues and 4 traits). Interestingly, BMD’s G×S signals are very strongly enriched in eQTL with associated eGenes (including *ABO* and *CYP3A5*) expressed in the adrenal gland, among other tissues. For height, we observed small enrichment *p*-values across many tissues since G×S focal SNPs are enriched in eQTL with associated eGenes (including *LOC101927975* and *CNDP2*) expressed across many tissues. Lastly for WHR, we observed G×S detections to be heavily enriched in eQTL with associated eGenes expressed in fibroblast, adipose, and skin tissues.

**Fig 5.**
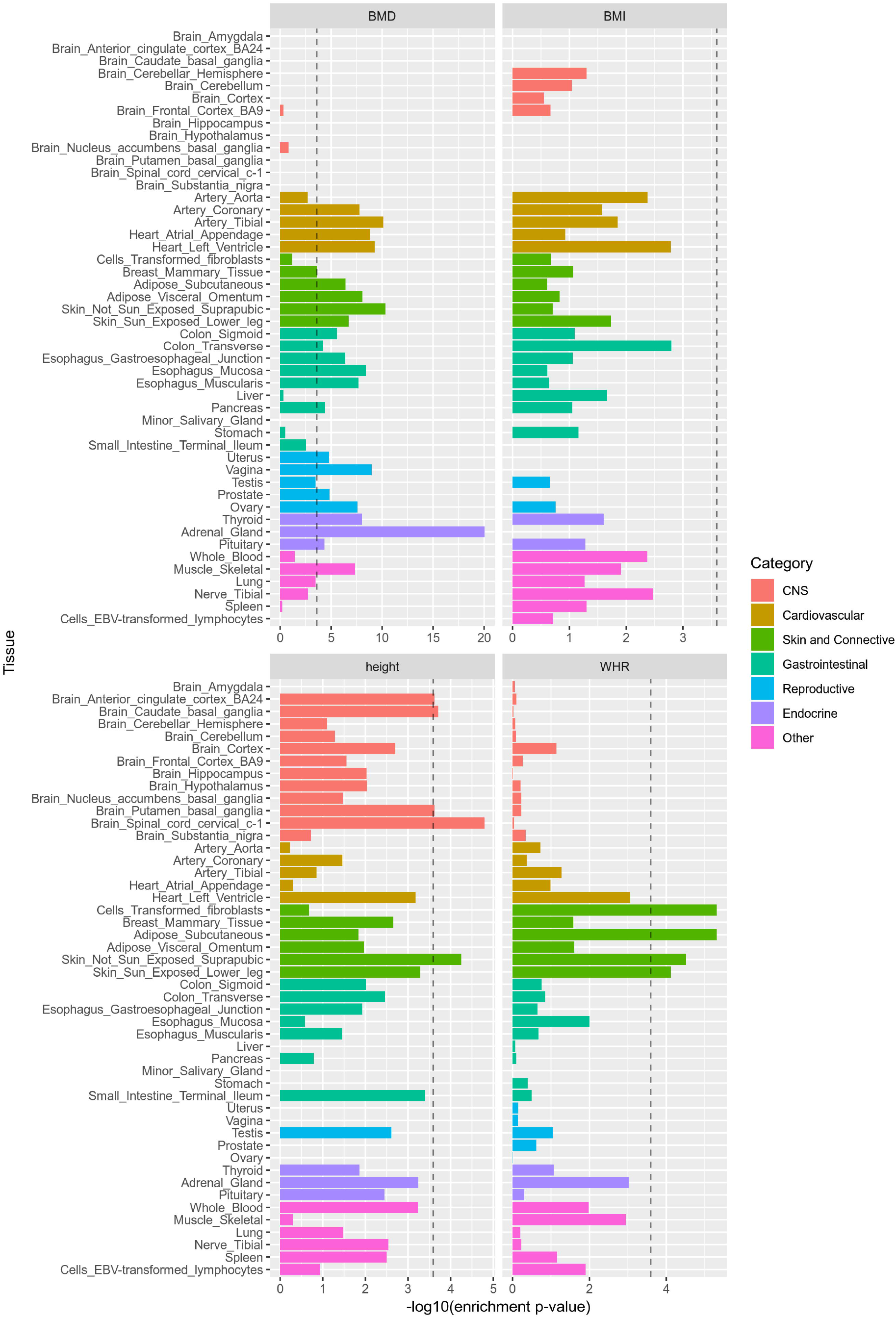
Evidence that LBR-identified G×S interactions are enriched in tissue-specific eQTL. Plotted on the x-axis is the *p*-value obtained from a hypergeometic test providing evidence that focal SNPs selected using 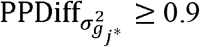 are enriched in tissue-specific eQTL. The dashed line represents a Bonferroni corrected significance threshold of 2.6×10^−4^.

## Discussion

We have investigated the degree to which sex-specific genetic architectures differ at local regions, using large biobank data (N ~ 119,000 males and ~140,000 females) and Bayesian multiple regression techniques that estimate sex-specific marker effects accounting for local LD patterns. The flexibility of the Bayesian approach enables multi-resolution inference of sex-specific effects: from individual SNP effects to window-variances that aggregate SNP effects within chromosome segments. These inferences can be drawn all using the results of the same model fit (eq. 1) but different post-processing of samples of SNP effects from the posterior distribution.

The Bayesian multiple regression technique performed in this study, along with estimation of window variances, was largely inspired by Fernando et al. [14]. In that study, windows were defined using disjoint, fixed intervals. In contrast, for each SNP we define a window based on local LD patterns, resulting in heavily overlapping, dynamically sized windows. The methods presented here also bear resemblance to those of Vilhjálmsson et al. [18], which utilized point-normal priors to estimate human SNP effects after accounting for local LD patterns. In that study, posterior means of SNP effects were estimated for the purposes of prediction while in this study, we numerically derive the full posterior distribution, allowing for inference of non-null SNP effects and window variances.

Through simulations, we showed that local Bayesian regressions (LBR) provide superior power and precision to detect causal variants and those specifically bearing G×S interactions. We rationalize improvements in power upon traditional SMR methods by noting that the magnitude of a typical causal variant or G×S interaction is exceedingly small and can elude hypothesis testing partly due to the burden of multiple test correction. We also note that the resolution (peak size) in SMR signals is relatively large when using large sample sizes (due to not fully accounting for local LD patterns). To overcome this problem, we provided evidence that LBR methods—either by estimating true marker effects or by aggregating true marker effects within relatively small regions—can achieve improved resolution when working with large sample sizes such as biobank-level data.

When using LBR to analyze real human traits, we have provided credence to our posterior probability-based discoveries by determining that LBR-detected G×S interactions are generally more enriched in eQTL than SMR-detected interactions. For BMD, we provided new evidence that sex-specific effects differ within *ABO* and that G×S interactions are highly enriched in adrenal gland-specific eQTL. This encourages the hypothesis that some G×S are eQTL that may modulate gene expression in the adrenal gland, with gene function dependent on the presence or absence of sex hormones. This was also an intriguing finding given that ABO blood groups have been known to associate with osteoporosis and osteoporosis severity [34,35]. For WHR, we detected previously known, large-magnitude G×S interactions that were discovered using WHR or WHRadjBMI [21–24], but additionally discovered novel, small magnitude G×S interactions near such genes as *ESR1* and *ABCA1.* In a previous work analyzing WHRadjBMI, *ABCA1* showed a significant female-specific genetic effect only, however the test for G×S interaction failed to reach significance [24].

For traits like height and BMI, large effect loci are estimated to have very similar effects between males and females and loci with evidence of G×S interactions were those possessing relatively small sex-specific effects. As seen in Fig 4B, many relatively large window variances for height are estimated to be slightly higher for males than for females albeit not reaching a 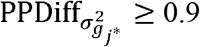 threshold. This is consistent with the fact that the global genomic variance for height was estimated to be higher in males than in females in a previous study using the interim release of the UK Biobank [4]. Similarly, the same prior study estimated the global genomic variance of BMI to be higher in females than in males and we observe, if anything, evidence of sex-specific window variances leading to the same conclusion. These observations may potentially indicate that relatively large causal variants have slightly different sex-specific effects for traits like height and BMI, however, if that is the case we are still underpowered to confidently detect such interactions.

It is important to acknowledge that while the methods presented here appear useful to decipher sex-specific genetic architectures from large human samples, additional work will be required to determine how these techniques may infer heterogeneous genetic effects in other contexts (other types of gene-by-covariate interactions), or when using different sample sizes or samples from different populations. With large sample sizes, the increased power and flexibility of the LBR comes with the cost of a significantly larger computational burden than the one involved in the traditional SMR approach; however, working with large datasets can be made manageable by adjusting size of each fitted segment (Fig 1) and parallel processing the fitting of each segment. Alternatively, LBR may be used as a follow up to traditional SMR tests, using pre-selected regions of interest. Another limitation inherent to aggregating SNP effects using window variances is that the sign of the effect is lost. In this way, when inferring G×S interactions through window variance differences, we cannot comment on whether sex-specific effects had the same sign or differing signs.

To conclude, we have demonstrated the powerful and flexible use of local Bayesian regressions for GWA to infer sex-specific genetic effects and G×S interactions using the UK Biobank. This was largely done by showing various means to utilize estimates of true (accounting for local LD), sex-specific SNP marker effects for GWA even when causal variants are not on the SNP panel for analysis. We anticipate that many more traits will be analyzed with this method to increasingly learn more about what is contributing to differences between males and females in human populations.

## Methods

### Genotype data

Individuals from the UK Biobank [36] were genotyped using the custom UK Biobank Axiom Array (http://www.ukbiobank.ac.uk/scientists-3/uk-biobank-axiom-array/) containing ~800,000 SNPs. SNP quality control proceeded with the Caucasian cohort (N = 409,700); SNPs with a minor allele frequency < 0.01 and missing call rate > 0.05 were removed. SNPs from sex chromosomes and the mitochondrial chromosome were not considered in this study, resulting in 607,497 autosomal SNPs. Individuals with coefficient of relatedness of 0.03 or greater were removed from analysis, resulting in 258,928 distantly related genotyped individuals for use in this study.

### Phenotype data

All phenotypic data was collected using baseline measurements of UK Biobank participants. For height, the description “Standing height” from the UK Biobank was used. Individuals with heights (cm) less than 147 or more than 210 were removed from analysis. For BMD, the descriptions “Heel bone mineral density (BMD)”, “Heel bone mineral density (BMD) (left)”, and “Heel bone mineral density (BMD) (right)” were used in conjunction; for individuals with missing “Heel bone mineral density (BMD)” records, either the (left), the (right), or if available, the average between (left) and (right) was used. For BMI, the description “Body mass index (BMI)” was used and for WHR, the ratio of “Waist circumference” to “Hip circumference” was used. Prior to model fitting, all traits were pre-corrected for sex, age, batch, genotyping center, and the first 5 principle components derived from genomic data. The adjusted phenotypes consisted of least-squares residuals from a model that included the effects listed above. For each trait, sample sizes and within-sex summary statistics are provided in S1 Table.

### LBR hyperparameters

Hyperparameters used in the LBR model (eq. 1) were error variances for each sex, the proportion of nonzero effects for each SNP effect component, and the variances of nonzero effects for each SNP effect component 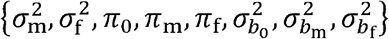. Variances (of either SNP effect components or sex-specific errors) were given a scaled-inverse Chi-square prior, parameterized by a degree of freedom parameter *df* (set to 5) and scaling parameter *S. S* is set according to built-in rules of the BGLR package using a prior model R-squared of 0.03 for main effects and 0.01 for the sex-interaction terms. More detail on how the scale parameter *S* is calculated can be found in Perez and de los Campos, 2014 [27]. *π_k_* was given a beta prior with shape parameters *α* = 2 and *β* = 2. An example of how to implement LBR (eq. 1) using BGLR with the above hyperparameter specifications is provided at https://github.com/funkhou9/LBR-sex-interactions.

### Inference using post-processing of posterior samples

BGLR uses Markov chain Monte Carlo (MCMC) to sample from the posterior distribution of sex-specific effects. For each MCMC sample we derived male and female effects using 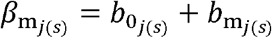 and 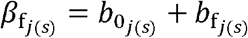, where *s* = 1,…, 4,350 indexes MCMC samples. Here, results were obtained using three separate MCMC chains. Each chain was obtained using 3,400 MCMC samples; the first 500 samples were discarded as burn-in and the remaining samples where thinned by an interval of 2, leading to 1,450 samples per chain.

Estimates of sex-specific SNP effects (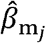 and 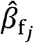) were obtained from their posterior means. We estimated the posterior probability of a female-specific non-zero SNP effect using 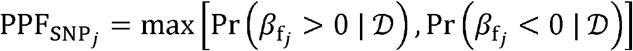, where 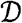 represents the observed data. This was done by counting the proportion of *β*_f_*j*__ samples above zero and below zero. This was repeated for inferring the male-specific SNP effect. The posterior probability of sex-difference at individual SNP-effects was estimated using 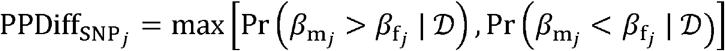 where again these probabilities were estimated using the corresponding frequencies from the posterior distribution samples.

For each MCMC sample we also aggregated SNP effects within window *j*\* using 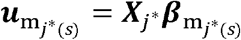 and 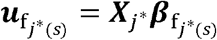. For this calculation we used a common genotype matrix ***X***_*j*\*_ consisting of all *N* male and female genotypes to avoid differences in additive genetic values arising from allele frequency differences between males and females occurring by random sampling. Samples of sex-specific window variances were obtained using the sample variance: 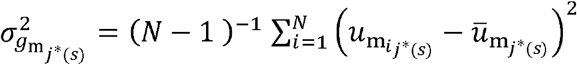 and 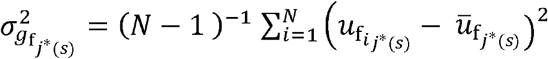. Estimates of sex-specific window variances were obtained from their posterior means. Inferring sex-specific window variances was done by estimating 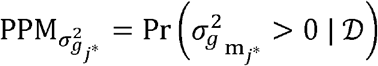 and 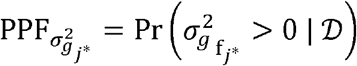 and inferring a G×S interaction at window *j*\* was done by estimating:

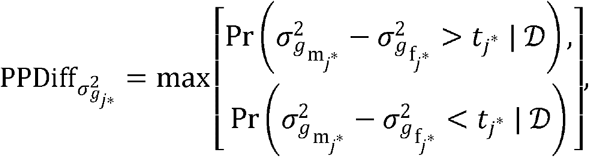

where *t_j*_* was used to exert judgment about how different sex-specific window variances must be to declare a meaningful G×S interaction. Here, *t_j*_* was one-tenth of the mean of all posterior samples of 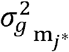 and 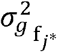. Functions to process posterior samples to estimate and infer non-null sex-specific effects and G×S interactions is provided at https://github.com/funkhou9/LBR-sex-interactions.

### Defining local, LD-based windows

To define SNPs contained within window *j*\*, a region of LD centered on SNP *j*, we collected all SNP *j*′ immediately surrounding SNP *j* for which *Cor*(***x***_*j*_,***x***_*j*\*_)^2^ ≥ 0.1. We allowed up to two consecutive SNPs in which *Cor*(***x***_*j*_,***x***_*j*′_)^2^ < 0.1 to allow for potential mapping errors or other unexplained instances where LD with SNP *j* dips only briefly. The function `getWindows()`, which provides windows given a genotype matrix ***X***, is provided in https://github.com/funkhou9/LBR-sex-interactions.

### Single marker regression

We also performed single-marker regression analyses using following model:

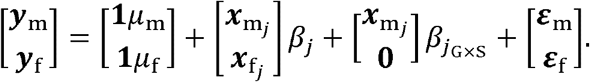

As with the LBR model (eq. 1), we assume sex-specific errors are distributed normally with zero mean and sex-specific variances. SNP effects and interactions were estimated using weighted least squares. To test for a G×S interaction at SNP *j*, a t-test is used: 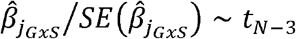. The *p*-value from such a test is referred to as *pvalue*-diff. To test for any association (either among males, females, or both), we used an F-test, comparing a restricted model: 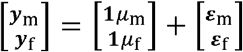 against the unrestricted model: 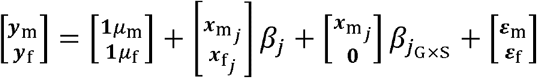.

### Simulations

Simulated traits were developed using 60,000 genotyped SNPs (the first 6,000 SNPs from the first ten chromosomes) from 119,190 males and 139,738 females. Using these SNP genotypes, each trait was simulated as follows:

1. A total of 150 causal variants (CVs) were randomly sampled from 60,000 SNPs.

- Let 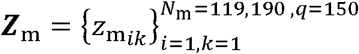 and 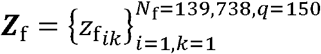 denote matrices of male and female genotypes at sampled CVs.
2. Additive CV effect sizes were randomly sampled from the gamma distribution. 90 CVs (those with homogenous effects) were sampled from Gamma(*k* = 10, *θ* = 1) and were made negative with a probability of 0.5. Of the 60 CVs with differing sex-specific effects, 30 had nonzero effects in both sexes but with deferring magnitudes: at random one sex’s effects were sampled from Gamma(*k* = 5, *θ* = 1) and the other from Gamma(*k* = 20, *θ* = 1). For the remaining 30 CVs, at random one sex’s effects were exactly zero while the other sex’s effects were sampled from Gamma(*k* = 10, *θ* = 1).

- Let 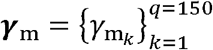 and 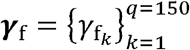 denote vectors of male-specific and female-specific CV effects, respectively, for all 150 CVs.
3. Error variances for males 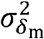 and females 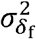 were adjusted such that the proportion of phenotypic variance explained by all QTL is 0.05 for both males and females (on the complete genome scale this corresponds to a heritability of about 0.5).

- Let 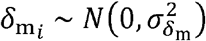 and 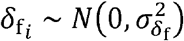 denote residual error for the *i*^th^ male and *i*^th^ female.
4. Male traits 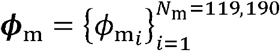 and female traits 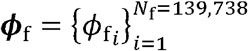 were simulated from a linear combination of QTL genotypes plus a residual error:

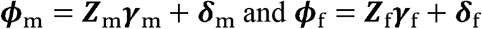
5. Steps 1-4 are repeated for 30 Monte Carlo replicates.

## Acknowledgments

Enrichment analysis performed in this manuscript was done using data from the Genotype-Tissue Expression (GTEx) Project. Single-tissue cis-eQTL data was downloaded from https://gtexportal.org/home/datasets on 02/01/19.

## Supporting information

**S1 Fig. LD statistics across distances.**

**S2 Fig. Estimated power and false-discovery rate for discovering observed SNPs with effects in at least one sex.** Estimated power (left) and FDR (right) shown as a function of the number of SNPs selected. Each point represents a sample average and error bars represent 95% confidence intervals, each derived using 30 Monte Carlo replicates. LBR (SNP): local Bayesian regression, utilizing PP_SNP_*j*__. SMR: single-marker regression, utilizing the F-test-based *p*-value.

**S3 Fig. Power vs false-discovery rate for discovering genomic regions containing masked causal variants.** Here power is defined as the expected proportion of causal variants that are being tagged by at least one selected SNP *j* or window *j*\*. False discovery rate is defined as the proportion of selected SNPs or windows that are not tagging any causal variants. Each point is an estimate and error bars for both axes represent 95% confidence intervals. Point estimates and intervals were derived using 30 Monte Carlo replicates. Each facet corresponds to a different “target area”, a fixed width around each causal variant that defines the set of SNPs effectively tagging it. LBR (SNP): uses the PP_SNP_*j*__ metric spanning 1-0. LBR (Window): uses the maximum between 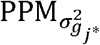 and 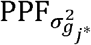 spanning 1-0. SMR: uses the F-test-based *p*-value spanning (on the −log_10_ scale) 30-0.

**S4 Fig. Comparison between SMR and LBR for discovering G×S interactions.** Manhattan plot showing *pvalue*-diff for each analyzed SNP. SNPs are colored yellow if they were focal SNPs with a 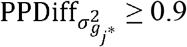 and colored red if they were focal SNPs with a 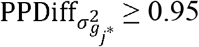. The dashed horizontal lines denote *p*-diff thresholds of 1×10^−5^ and 5×10^−8^.

**S5 Fig. eQTL enrichment as a function of the number of SNPs selected.** LBR (Window): uses the 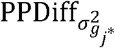 metric. SMR: uses the *pvalue*-diff metric.

**S1 Table. Sex-specific phenotype statistics.** Height units: cm, BMD units: g/cm2, BMI units: Kg/m2.

**S2 Table. Inferred G×S interactions using sex-specific window variances.** Listed are all windows with a 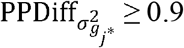.

