## Supplementary figures and images for "Deciphering sex-specific genetic architectures using local Bayesian regressions"

### Supplemental Figure 1

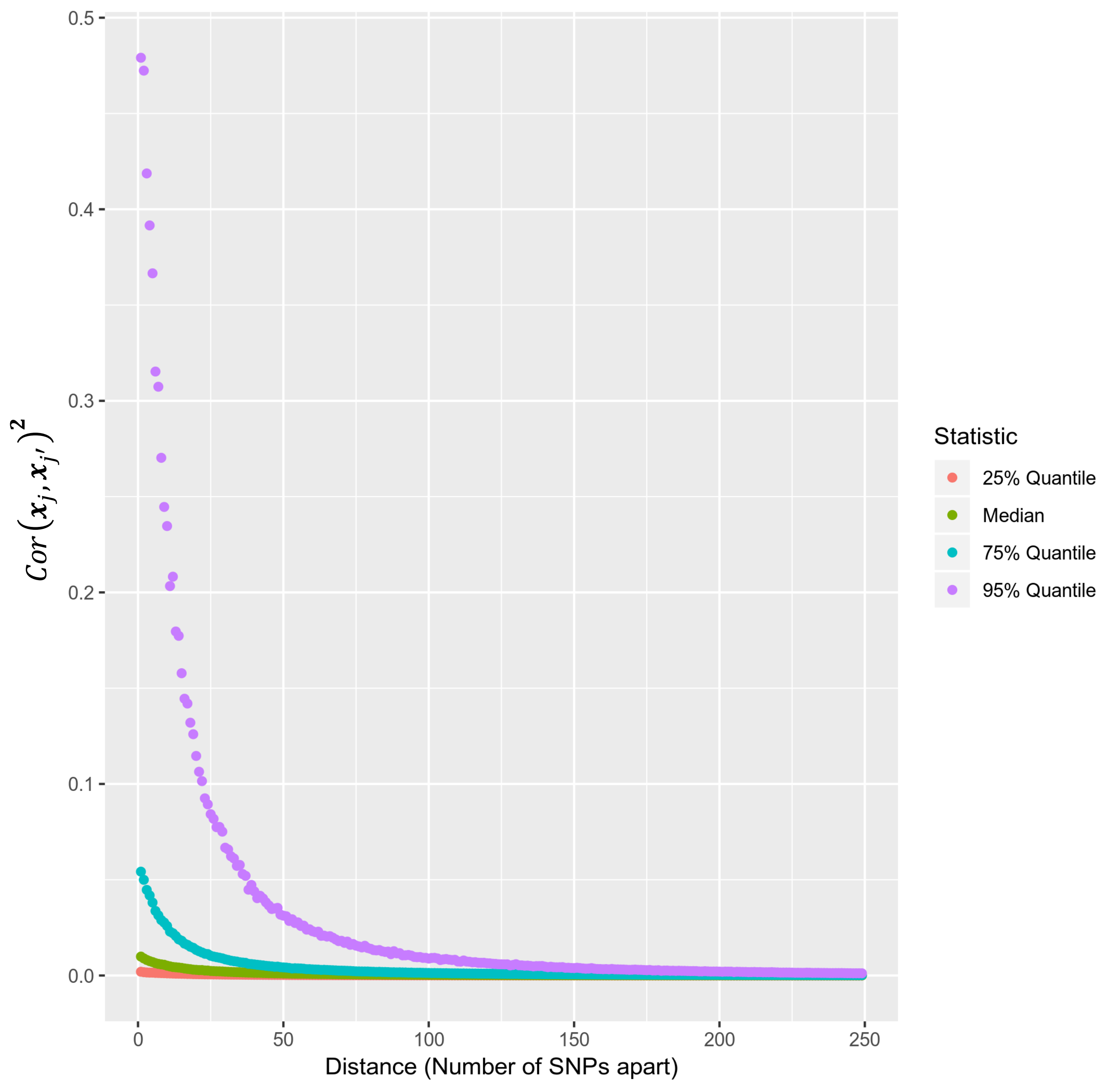

### Supplemental Figure 3

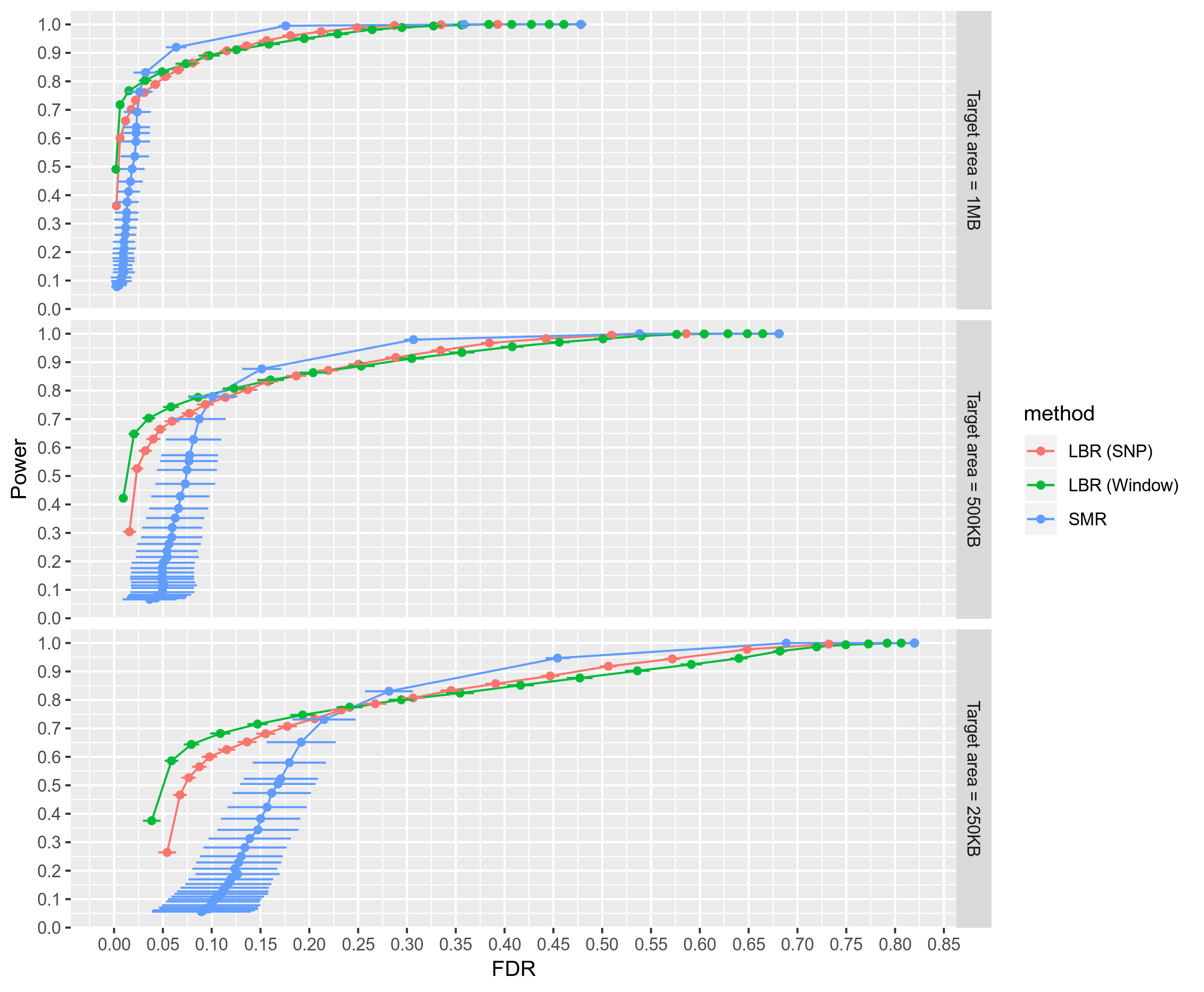

### Supplemental Figure 4

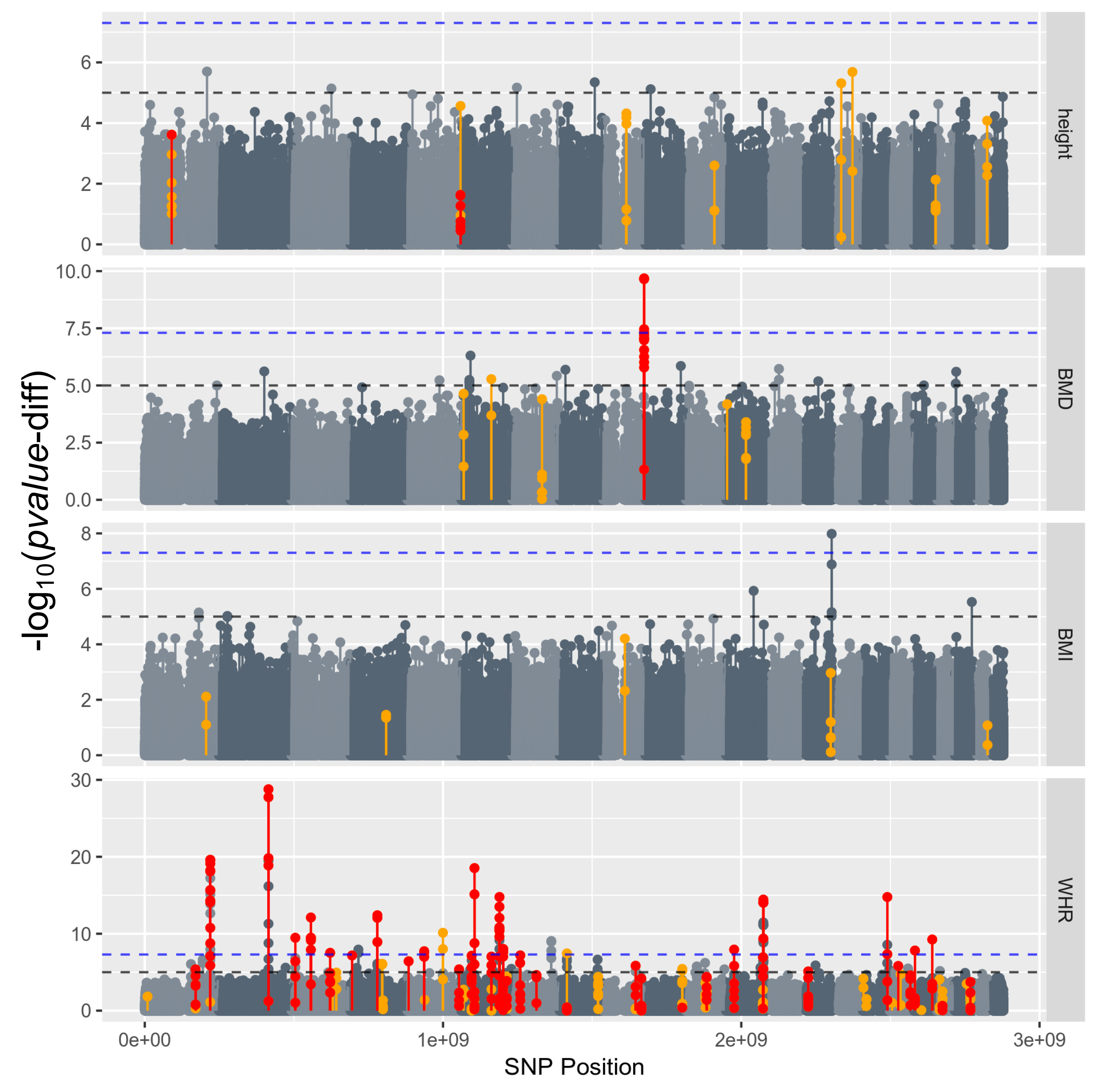

### Supplemental Figure 5

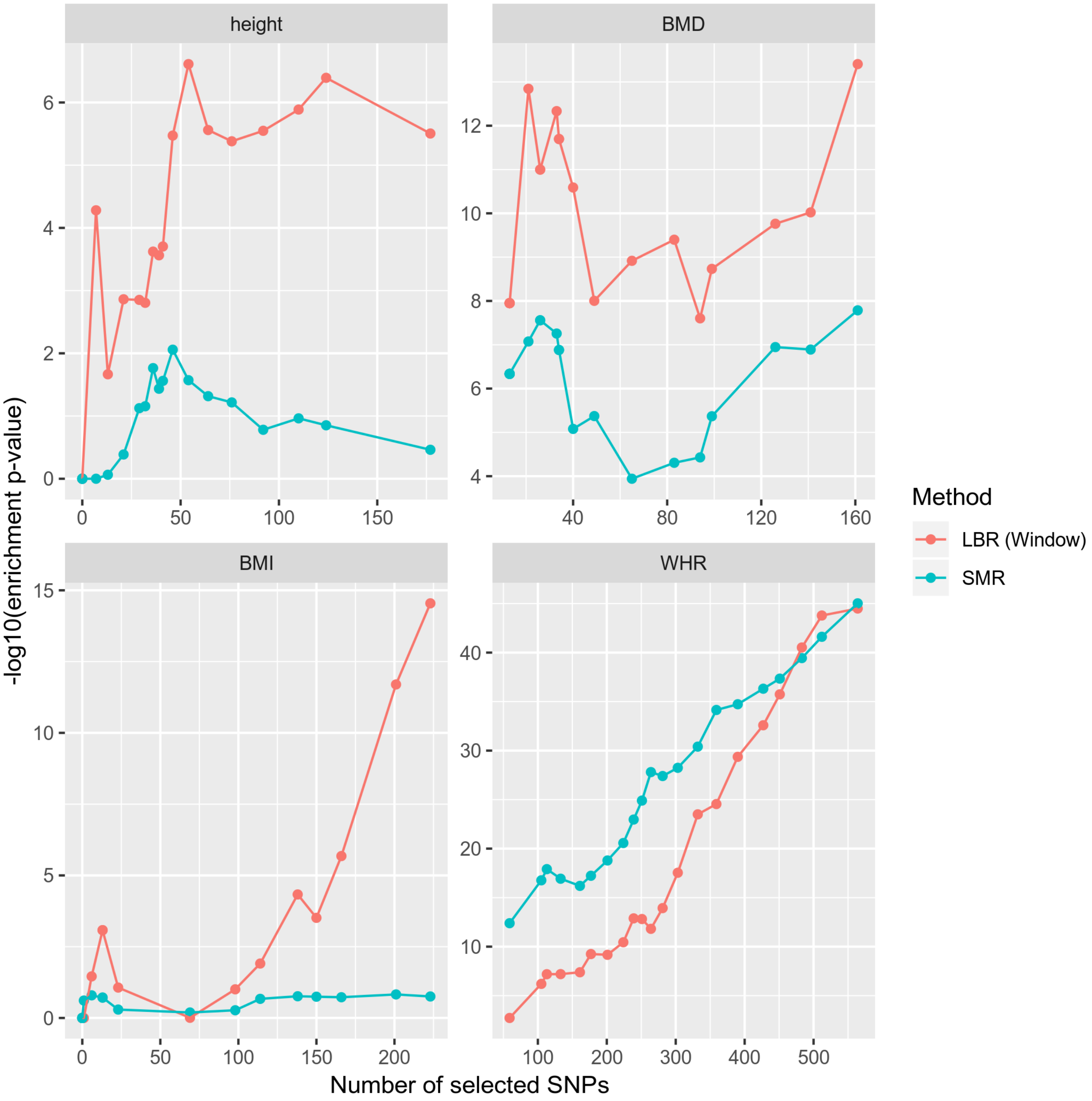

### Supplmental Figure 2

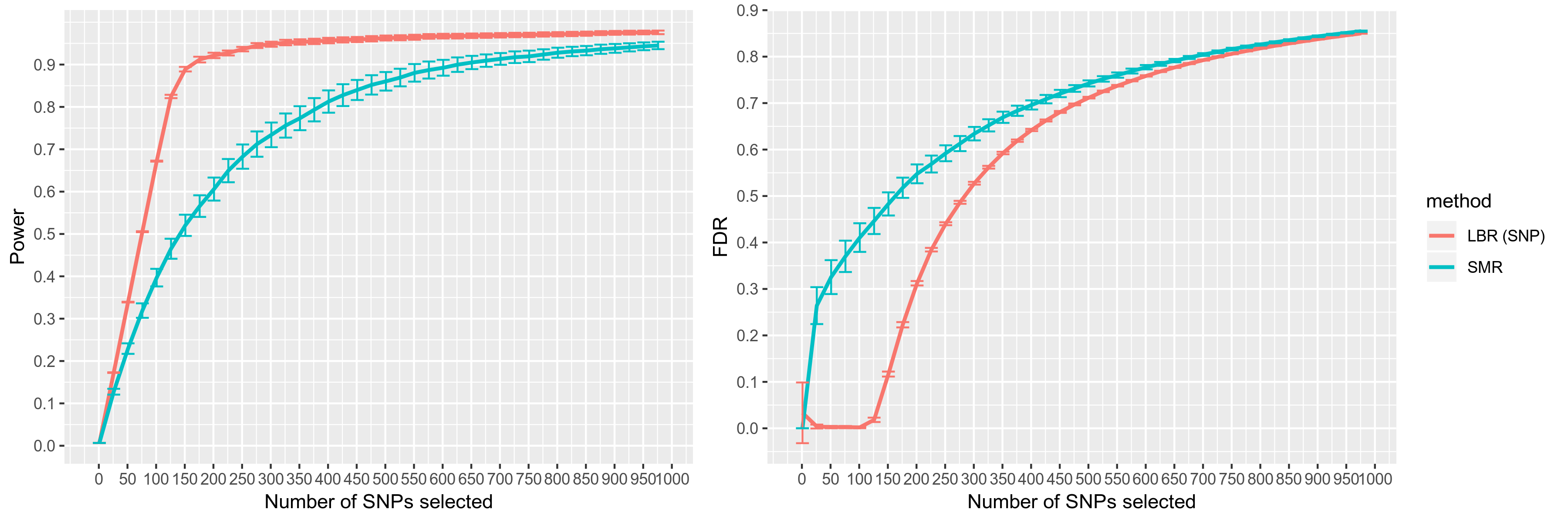
